# Romiumeter: An Open-Source Inertial Measurement Unit-Based Goniometer for Range of Motion Measurements

**DOI:** 10.1101/2024.05.29.596353

**Authors:** Basinepalli Kothireddy Gari Diwakarreddy, S Abishek, Amal Andrews, Lenny Vasanthan, Sivakumar Balasubramanian

## Abstract

Range of motion (ROM) serves as a crucial metric for assessing movement impairments. Traditionally, clinicians use goniometers to measure the ROM, but this method relies on the clinician’s skill, in particular for difficult joints such as the shoulder and neck joints. Recent studies have explored the use of wearable inertial measurement units (IMUs) as an alternative. IMUs exhibit excellent agreement with goniometers, but the lack of affordable, accessible, and clinically validated tools remains an issue. This paper introduces the Romiumeter, a single IMU-based device designed to measure the ROM of the neck and shoulder movements. To validate its accuracy, the Romiumeter was tested on 34 asymptomatic individuals for shoulder and neck movements, using an optical motion capture system as the ground truth. The device demonstrated good accuracy, with a maximum absolute error of less than 5*°* with moderate to good reliable measurements(inter-rater reliability: 0.69 - 0.87 and intra-rater reliability: 0.76 - 0.87). Additionally, the Romiumeter underwent validation for different algorithms, including the complementary and Madgwick filters. Interestingly, no significant differences were found between the algorithms. Overall, the Romiumeter provides reliable measurements for assessing shoulder and neck ROM in asymptomatic individuals.

## 1 INTRODUCTION

Objective measures of human movement ability are essential for the diagnosis/prognosis of various neuro-musculoskeletal conditions [1]. Amongst such measures, a joint’s range of motion (ROM) is a fundamental measure evaluating movement impairments, which is a basic marker for many impairment guides and clinical outcome studies [2, 3]. Thus, a valid, reliable, and sensitive measurement of ROM is a vital component in a clinician or researcher’s toolbox for assessing movement behaviour.

The gold standard approach for measuring the ROM is the universal goniometer – a highly practical, compact, portable, and easy-to-use tool [4]. It comprises two arms hinged at one end with a 360*°* protractor. Measuring the ROM using the goniometer involves stabilizing one arm at the proximal part of the joint while manoeuvring the other arm with the joint’s distal part to determine the motion’s arc across the joint [1]. The resulting joint angle is then easily read out from the protractor. Although a simple and highly usable tool, its accuracy is impacted by several factors such as the tester’s skill, misalignment between the joint and the goniometer, misidentification of standard bony landmarks [5], and difficulties in measuring the ROM of specific joints (e.g., neck joint). Additionally, the universal goniometer’s inter-rater reliability has a large variation across studies (ICC range: 0.25-0.91) compared to intra-rater reliability (ICC range: 0.81-0.94) [6–8]. Boone et al. reported that the inter-rater reliability was higher for the upper than for the lower limb [9]. Hence, there is a need for a better tool for measuring the ROM that performs at least as well as the standard goniometer and is robust to the tester’s skills and the specific joint being assessed. This tool should be compact, portable, easy to use, and reliable.

Several other technologies have been explored in the literature to address this limitation. These include digital inclinometers [10–12], electromagnetic motion analysis systems [13–16], and CROM (Cervial Range-of-Motion) based on the gravity plus compass goniometry) [17], which were tested for their validity and reliability in measuring the ROM for different joints. Although digital inclinometers are simple and reliable, they require the participant to be positioned in the appropriate orientation with respect to gravity to measure the ROM for different joints. Furthermore, to the best of our knowledge, there are no studies that validate their accuracy against highly accurate systems such as Optical Motion Capture System (OMCS). Electromagnetic motion analysis systems like FASTRAK and Flock of Birds rely on the electromagnetic field. These can only be used in environments free of magnetic/metallic substances and require extensive calibration [18]. The CROM has demonstrated good reliability [19] and an excellent linear relationship with the OMCS. However, it is designed for the cervical joint and requires an environment without any strong magnetic disturbances to measure the ROM for left/right rotational movements.

Several studies have explored ultrasound and image processing-based methods for measuring the ROM [20–23]. The OMCS was found to be the most accurate and is considered the gold standard for measuring movement kinematics [24]. However, the OMCS is not practical for routine clinical use for single joint ROM measurement due to its setup time, cost, etc. In recent years, there has been an increasing interest in motion-tracking using wearable inertial measurement units (IMU), such as the 6 degrees-of-freedom (DOF) IMU with an accelerometer and a gyroscope [25–27]. The linear acceleration and the angular velocity of a limb measured by the IMU are used to estimate the orientation of the limb, which is used for computing a joint’s ROM. Researchers have explored their validity and reliability across various contexts, including gait analysis, joint range of motion assessment, and post-surgical monitoring [28–30]. There is an excellent agreement in the measurements in a comparison study between the goniometer and IMU sensor for measuring the ROM of the shoulder joint when assessed by a single tester (ICC > 0.90) [25]. However, there is a lack of availability of affordable, accessible, and clinically validated IMU-based tools for measuring the ROM that a clinician can employ in daily practice.

The current work aims to address this need by developing and validating an open-source single IMU-based device and the associated PC-based software for assessing the ROM of the neck and shoulder joints in all three axes. To this end, this work focuses on:

1. developing a single IMU-based, compact, battery-operated ROM measurement device.
2. validating the sensor’s measurements against an OMCS in asymptomatic individuals and
3. Measuring the intra-rater and inter-rater reliability of the sensor and its protocol for assessing neck and shoulder ROM in asymptomatic individuals.

## 2 Methods

The study aimed to develop and validate an open-source IMU-based device for measuring the ROM of the different human joints; we focused on the shoulder and neck joints in the present study. Designed to work like an electronic goniometer for measuring ROM, we call this device the *Romiumeter*.

### 2.1 Romiumeter: Hardware and Firmware

The Romiumeter was designed to have a compact form factor that can be worn on any body segment using a Velcro strap. The current version was built using off-the-shelf components, which included a Seeeduino XIAO board with ARM® Cortex®-M0+ 32bit 48MHz microcontroller (SAMD21G18), a 6-DOF IMU sensor (MPU6050, InvenSense-TDK Co.), an HC05 Bluetooth module, and a TP4056 1A Li-ion lithium battery charging module. The device was powered by a 600mAh lithium polymer battery, which can run for 5-6 hours on a full charge. The exploded 3D model view of the Romiumeter is shown in Fig. 1(A). This enclosure was 3D printed using a Flashforge Guider 2S printer, and Velcro straps were attached to the 3D-printed side loops. A picture of a participant wearing the Romiumeter on the upper arm is shown in Fig. 1(B). The Romiumeter’s firmware was developed using the Arduino IDE, and it was programmed to run at 588Hz (the maximum sampling frequency that was achieved given the constraints of the Bluetooth module) to read and transmit the accelerometer and gyroscope data (16-bit resolution) via the Bluetooth serial protocol to a PC. A custom communication protocol was designed to transmit this multi-byte data as individual packets of size 16 bytes (refer to the appendix A.1 for details).

**FGURE 1.**
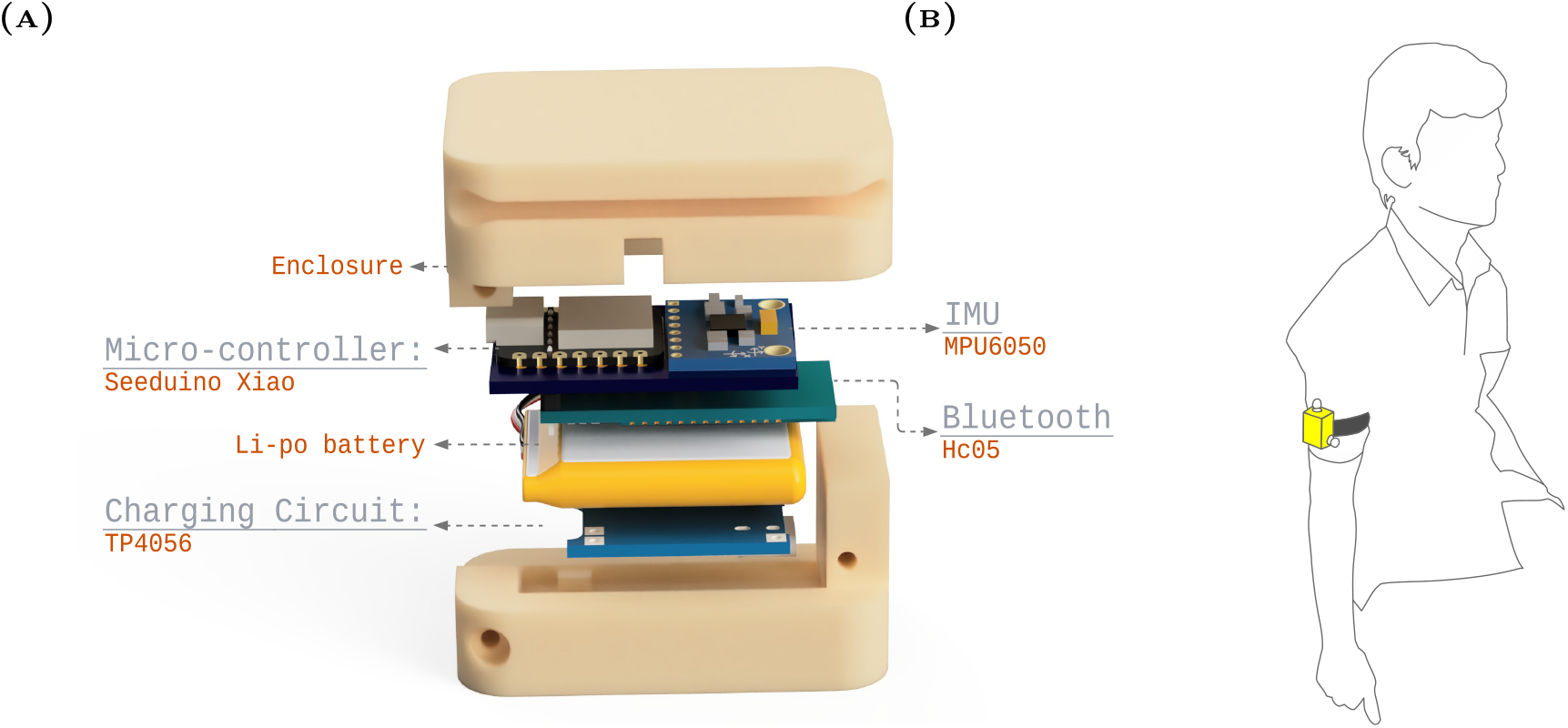
A) Exploded view of the Romiumeter B) A participant wearing the sensor on his upper arm.

### 2.2 Validation of Shoulder and Neck ROM Measurements with the Romiumeter

To develop the Romiumeter into a tool for measuring joint ROM, we validated the accuracy and reliability of its ROM measurements by comparing them against the measurements from an OMCS. This was performed on asymptomatic individuals performing different neck and shoulder joint movements using a custom assessment protocol. The validation study was approved by the Christian Medical College (CMC) Vellore’s Institutional Review Board (IRB Min. No.14668 dated 08/06/22).

Asymptomatic participants donned the Romiumeter on their upper arm/forearm or the forehead to measure the ROM of the shoulder or neck joint, respectively. For shoulder ROM measurements, participants performed multiple repetitions of three types of shoulder movements: shoulder flexion/extension, shoulder abduction/adduction, and shoulder internal/external rotations. For neck ROM measurements, participants performed multiple repetitions of neck flexion/extension, neck left/right lateral flexion, and neck right/left rotation. The cartoons of these different shoulder and neck movements are depicted in Fig. 2. The participants repeated these movements three times, using the following protocol for each repetition:

**FGURE 2.**
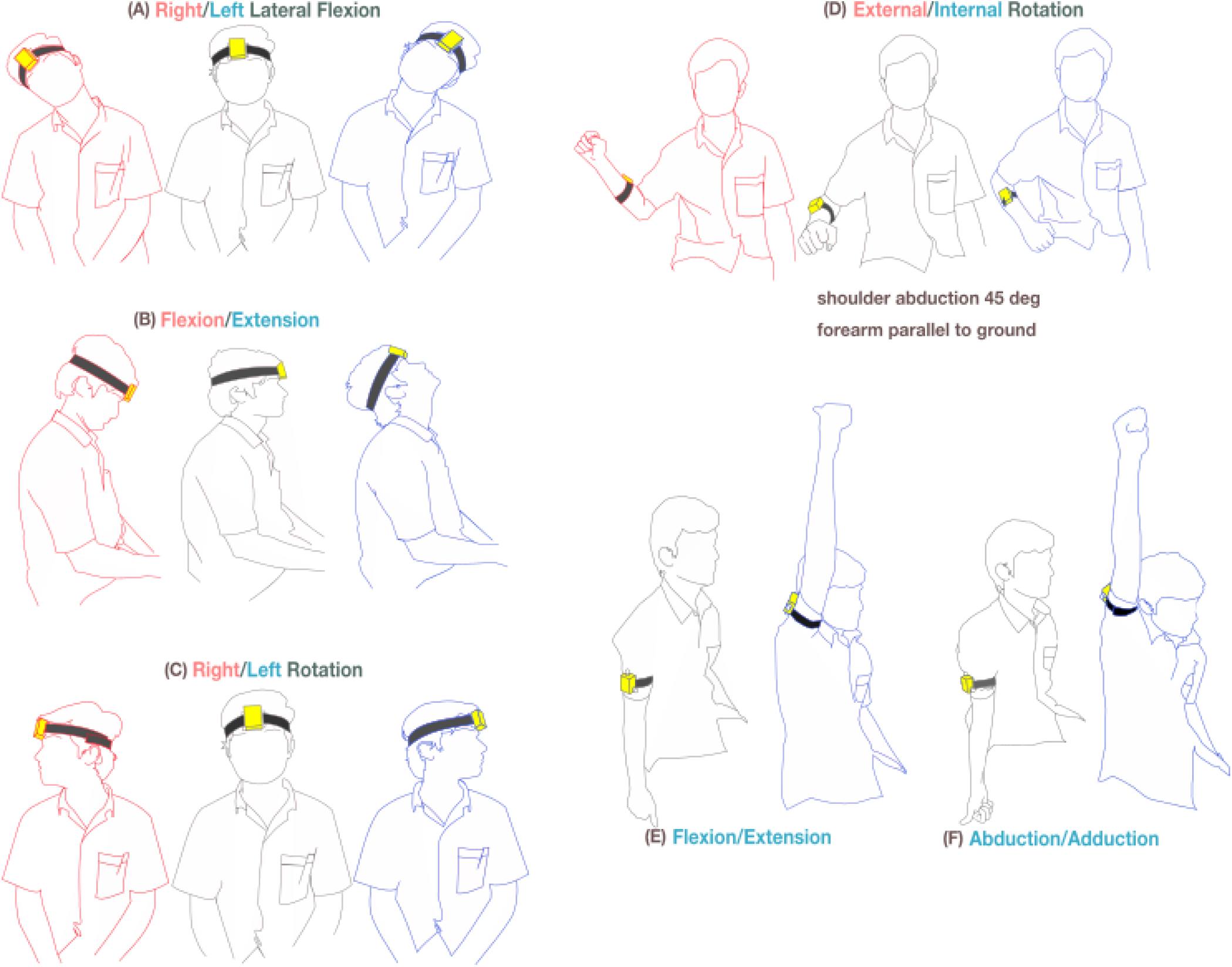
The illustrations above depict various shoulder and neck movements conducted in the study. The left column illustrates neck movements. A) represents the right lateral flexion in red, the left lateral flexion in blue, and the resting position in grey. B) represents the flexion in red, the resting position in grey, and the extension in blue. C) represents the right rotation in red, the resting position in grey, and the left rotation in blue. The right column represents shoulder movements. D) represents the external rotation in red, the resting position in grey, and the internal rotation in blue. E) and F) represent the flexion/extension and the abduction/adduction movements in blue, with the resting position depicted in grey.

1. Participants rested (no movement) for 2-4 seconds in the resting posture defined for each movement type.
2. Upon the “go” instruction from the experimenter, the participants performed a single repetition of a particular movement and returned back to the resting posture. The participants were requested to move at a comfortable speed to the maximum ROM they could, neither too fast nor too slow.

The experimenter, a trained bachelor of physiotherapy student conducting the assessment, ensured, through visual inspection, that the participants (a) started from the appropriate resting posture without any visible movement during the rest phase and (b) performed the appropriate movement without interfering with the movements of the trunk or other joints. If these conditions were violated, the participant was given feedback, and the trial was repeated.

#### 2.2.1 Participants

The validation of the Romiumeter was carried out on asymptomatic participants. The inclusion criteria for the asymptomatic volunteers were: (i) aged 18 years and above, (ii) no history of shoulder and neck pain in the past 3 months, (iii) no movement restriction in the shoulder and neck joints, and (iv) willing to attend three measurement sessions on two different days within a week’s gap.

The participants attended three measurement sessions over two days. On each session, the participant donned the Romiumeter on their upper/forearm and forehead with a set of reflective markers placed on the Romiumeter’s enclosure. Two physiotherapists with adequate experience in ROM measurement acted as the experimenters to collect data from all asymptomatic participants. To check the validity and reliability of Romiumeter, on day 1, two measurement sessions were carried out with the participant, with a gap of 1 hour. The same physiotherapist administered both these measurement sessions. On day 2, the other physiotherapist did the third measurement session.

In each session, the participant donned the Romiumeter on the upper arm to perform the shoulder flexion, abduction/adduction and donned the device on the forearm to perform the shoulder internal/external rotation. Following this, the participant donned the Romiumeter on the forehead and performed neck flexion/extension, lateral flexion/extension, and right/left rotation. The sensor location on the body segment for the shoulder and neck joint assessment was standardized based on the following: (i) high sensitivity to the movement being performed, (ii) ease of donning, (iii) participant’s comfort, and (iv) visibility of the passive markers to the OMCS cameras. The data from these three sessions were used for evaluating (a) the accuracy of the joint ROM assessed using the Romiumeter by comparing it against the ground truth from the OMCS and (b) the intra- and inter-rater reliability of the ROM assessment protocol using the Romiumeter for the shoulder and the neck movements.

#### 2.2.2 Movement kinematic data recording

During each measurement session, the data from the Romiumeter were recorded on a PC running custom software presenting a graphical user interface. On another PC, the 3D positions of the infrared reflective markers on the Romiumeter were recorded using the OptiTrack motion capture system (NaturalPoint, Inc., Corvallis, OR, USA). A sync-out Transistor-Transistor Logic (TTL) pulse was sent out from the OptiTrack’s camera hub to a digital pin in the Romiumeter, which records the state of this digital pin along with the IMU data. The digital pin is set to 0 when the OptiTrack is not recording data and set to 1 during the recording of a trial. This digital pin’s state is used to synchronize the IMU and the OMCS data.

The OptiTrack system used in this study consists of 8 cameras that record the position of the reflective markers in a custom reference frame defined after calibrating the camera setup. The position data from the 3 reflective markers placed on the Romiumeter were logged at 100Hz.

### 2.3 Kinematic Reconstruction Algorithms

#### 2.3.1 From the OptiTrack Data

Fig. 3A shows the processing pipeline for extracting the angle of rotation of the shoulder/neck joint around its axis of rotation from the raw OCMS data. The 3D position of 3 markers was up-sampled through spline interpolation to match the sampling frequency of the Romiumeter (588Hz). A Savitzky-Golay filter [31] of window length 251 and order 3 (corresponds to the lowpass cut-off frequency of 6.0Hz [32]) was used to smoothen the position data from the three markers.

**FGURE 3.**
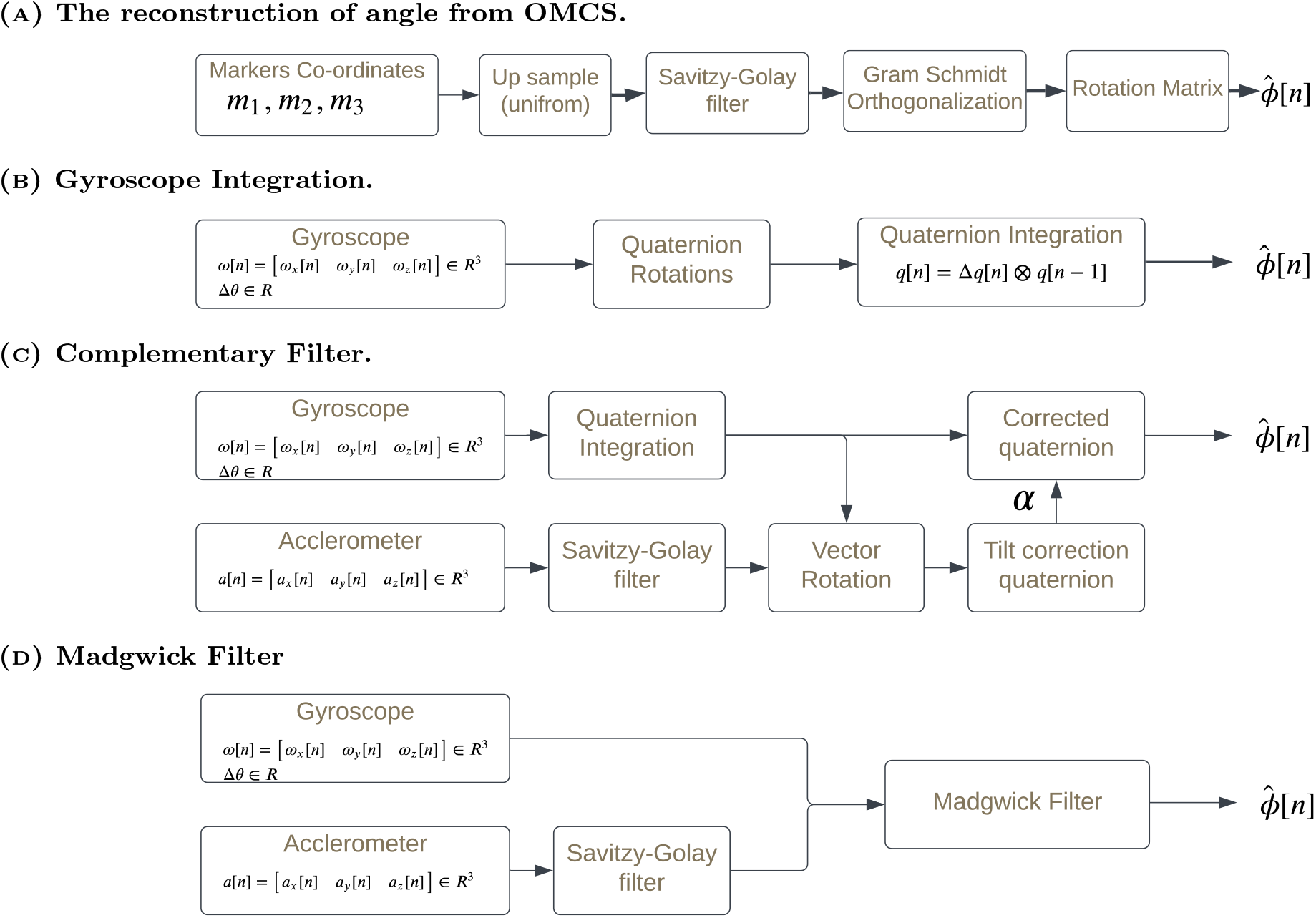

Let the position of the 3 markers with respect to the OMCS reference frame be **p**_1_ [*n*], **p**_2_ [*n*], **p**_3_ [*n*] ∈ ℝ^3^ at time instant *n* ∈ ℤ. Let’s define three vectors **m**_1_ [*n*], **m**_2_ [*n*], **m**_3_ [*n*] ∈ ℝ^3^, which together form a non-orthonormal reference frame rigidly attached to the Romiumeter.

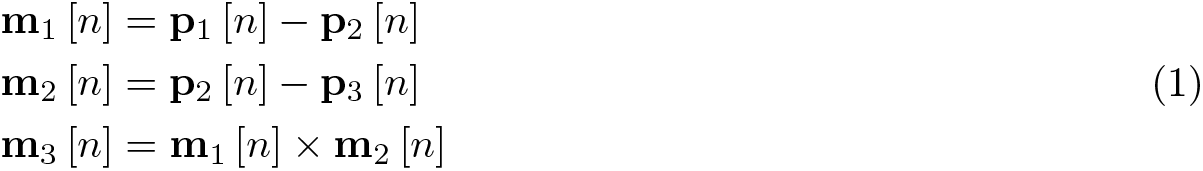

Applying, the Gram-Schmidt orthogonalisation procedure to the set {**m**_1_ [*n*], **m**_2_ [*n*], **m**_3_ [*n*]} results in an orthonormal set of vectors 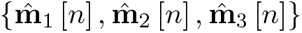. These vectors form an orthonormal reference frame attached to the Romiumeter, which can be represented using the rotation matrix 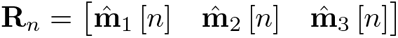 at time instant *n*. Assuming each of the three shoulder/neck movements of interest is a rotation about a fixed axis, we can obtain the angle of rotation of the joint from the rest position as follows,

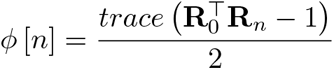

where, *ϕ* [*n*] is the angle of rotation at time instant *n* measured with respect to the orientation at time instant *n* = 0 (start of a trial).

#### 2.3.2 From the Romiumeter Data

Fig. 3B shows the processing pipeline for extracting the rotation angle of the shoulder/neck joint using three different approaches: (a) gyroscope integration, (b) a complementary filter [33], and (c) the Madgwick filter [34]. The gyroscope integration approach was used because of short-duration movements. The complementary and Madgwick filters were chosen because they are two popular methods for orientation reconstruction with IMUs.

##### Gyroscope integration

A gyroscope measures the angular velocity ***ω***[*n*] = [*ω*_*x*_ [*n*] *ω*_*y*_ [*n*] *ω*_*z*_ [*n*]] ∈ ℝ^3^ in its local sensor frame. For rotational movements about a fixed axis, the angle of rotation can b e obtained through quaternion integration to accumulate the rotation angle over time with respect to the initial reference frame at *n* = 0. However, it is well known that micro-electronic mechanical systems gyroscope signals suffer from a slow varying drift [35], which can lead to a wandering signal when integrated over a long duration. For relatively short-duration movement recordings, a basic model of a gyroscope signal can be given by the following:

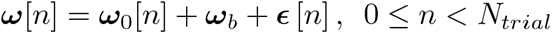

where, ***ω***[*n*] is the signal measured by the gyroscope at time instant *n*, ***ω***_0_[*n*] ∈ ℝ^3^ is the true angular velocity of the sensor in its local reference frame, ***ϵ*** [*n*] ∈ ℝ^3^ is the zero-mean random measurement noise, and ***ω***_*b*_ ∈ ℝ^3^ is the constant unknown offset error in the gyroscope signal. Because the offset is a slow-varying signal, it can safely be assumed to be constant over a short duration of interest 0 *≤ n < N*_*trial*_.

To minimize the effect of this offset ***ω***_*b*_ on the rotation angle estimate, an estimate of the offset is computed and subtracted from the gyroscope signal before performing the quaternion integration for each movement trial. The offset estimate is obtained through two approaches: (i) a one-time estimate 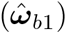 obtained by keeping the sensor at rest (***ω*** [*n*] = **0**) for 10 sec and computing the mean gyroscope signal over this period, and (ii) a trial-wise estimate 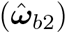 obtained from the first 500 samples of every movement trial when the sensor is at rest, i.e. ***ω***_0_ [*n*] = **0**, 0 *≤ n <* 500. The offset-removed gyroscope signal for each trial 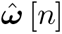, 0 *≤ n < N*_*trial*_ is computed by subtracting the trial-wise offset 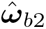 from the raw gyroscope signal, whenever the maximum standard deviation of the gyroscope signal across all three axes is within 1deg/sec, during the initial rest period of a trial. This threshold is defined as the value for which the average reconstruction error is minimum across all trials. If this condition is violated, i.e., if the body segment was not truly at rest during the trial, the one-time offset is used to remove the offset from the raw gyroscope signal. The details of the quaternion integration algorithm are described in appendix A.4.

##### Complementary Filter

The complementary filter combines the data from the accelerometer and gyroscope to provide an estimate of the angular kinematics of IMU [33]. This fusing of information from the accelerometer and the gyroscope can provide a better estimate of the angular kinematics of the IMU than simple quaternion integration of the offset-removed gyroscope signal. It should be noted that this can be done only for the pitch/roll and not yaw. The complementary filter implemented for this study is detailed in appendix A.5 and the calibration of accelerometer is detailed in appendix A.2.

##### Madgwick Filter

The Madgwick filter [34] is one of the most widely used sensor fusion algorithms for estimating an IMU’s angular kinematics, which employs gradient descent to correct for the gyroscope drift using the accelerometer data. Here also, the offset-removed gyroscope signal 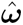 and the low pass filtered accelerometer signal are used as inputs to the filter. Refer to the appendix A.5 for the details of its implementation.

The gain parameter for the complementary and Madgwick filters was optimized using a 5 -fold validation method to minimize the angular reconstruction error with respect to the OMCS measurements. Refer to the appendix A.3 for more details. The results presented in this work are based on the optimal parameters estimated for the complementary and Madgwick filters.

### 2.4 Additional Experiments and Analysis

The previous validation experiment provided an estimate of the angle reconstruction (*ϵ*_*rec*_) and ROM (*ϵ*_*rom*_) error, which we quantified f or each movement trial as follows:

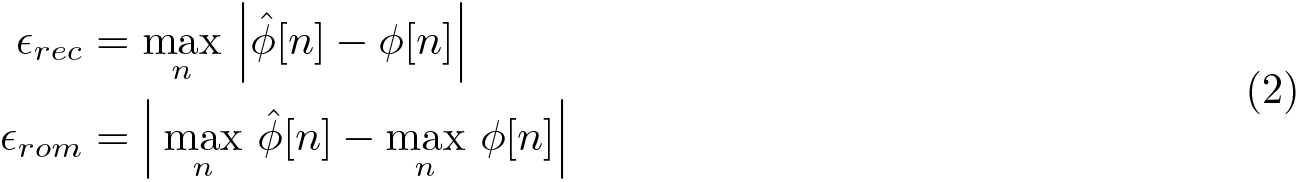

where 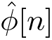 and *ϕ*[*n*] are the reconstructed angles from the Romiumeter and OMCS data, respectively, for a given trial. Note that the errors *ϵ*_*rec*_ and *ϵ*_*rom*_ provide an estimate of the worst-case errors due to the use of the max operator. There can be multiple sources for the reconstruction (and thus the ROM) error. The following additional experiments were performed to understand some of the potential sources of errors:

#### 1. Rest state reconstruction

One of the sources of error is the inherent random measurement noise in the IMU. To assess the impact of this noise on the overall error, we kept the Romiumeter at rest for 40 minutes and recorded the gyroscope data. This data was then divided into non-overlapping 10-second segments, resulting in 240 trials. The contribution of the noise to the observed error can be measured as the average of the maximum reconstructed angle over the trials; the reconstructed angle 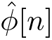 should be uniformly zero for all trials.

#### 2. Comparison with a high-resolution encoder

The accuracy of the OMCS system depends on several factors like the number of cameras, their relative locations, locations of the markers in the calibrated volume, and occlusions that result in the loss of data due to the loss of the sight of markers [36]. These factors could have contributed to the errors in the ground truth, contributing to the observed reconstruction errors. To determine the contribution of the OMCS system to the reconstruction error, we employed a high-resolution optical encoder (Encoder MILE, 6400 CPT, 2 Channels, with Line Driver RS 422, angular resolution = 0.01956*°*, Maxon Precision Motors Inc., Switzerland) to perform single DOF rotational movements of the Romiumeter (in different orientations). The movements of the Romiumeter were simultaneously recorded using the OMCS, as was done during the validation experiments with asymptomatic individuals. The setup joint could be placed in different orientations to simulate the movements of the different joints. Three movements were chosen for this experiment, which simulate: neck rotation, shoulder flexion/extension, and shoulder abduction/adduction. The protocol for each trial was the same as that for asymptomatic individuals except that the actual movement was done by rotating the encoder by hand.

### 2.5 Statistical analysis

The intra- and inter-rater reliability of shoulder and neck range of motion, as measured by the Romiumeter, were computed using a two-way random effects model, ICC(2,2), with a 95% confidence interval (CI). The ICC scores we interpreted as follows: scores less than 0.5 indicated poor reliability; scores between 0.5 and 0.75 indicated moderate reliability; scores between 0.75 and 0.9 indicated good reliability; and scores greater than 0.9 indicated excellent reliability. The intra-rater reliability was computed between the first two sessions conducted by the same physiotherapist on day 1. The inter-rater reliability was computed between the first session conducted by one physiotherapist on the first day and the third session conducted by a second physiotherapist on the following day. Pair-wise, one-way ANOVA was performed to check the statistical difference between the three algorithms used to compute the range of motion from the Romiumeter data. All the statistical analysis was performed in Python.

The full dataset used in this study and the code for the analysis are available at https://github.com/bkdiwakar34/Romiumeter.

## 3 Results

Thirty-four asymptomatic volunteers (19 males, 15 females; mean age of 22.70 years, SD 2.30) were recruited for the study through verbal advertisement in the Bioengineering and Physical Medicine and Rehabilitation departments at CMC Vellore. All participants signed an informed consent form before the recruitment, and no one was coerced to participate in the study. No financial compensation was provided for the participants. All participants attended three measurement sessions on two days. Fig. 4 shows a typical trial from the validation study from a participant performing shoulder external rotation movement. The plot displays the raw accelerometer and gyroscope data, along with the reconstructed angles from the OMCS and the three reconstruction algorithms using the Romiumeter.

**FGURE 4.**
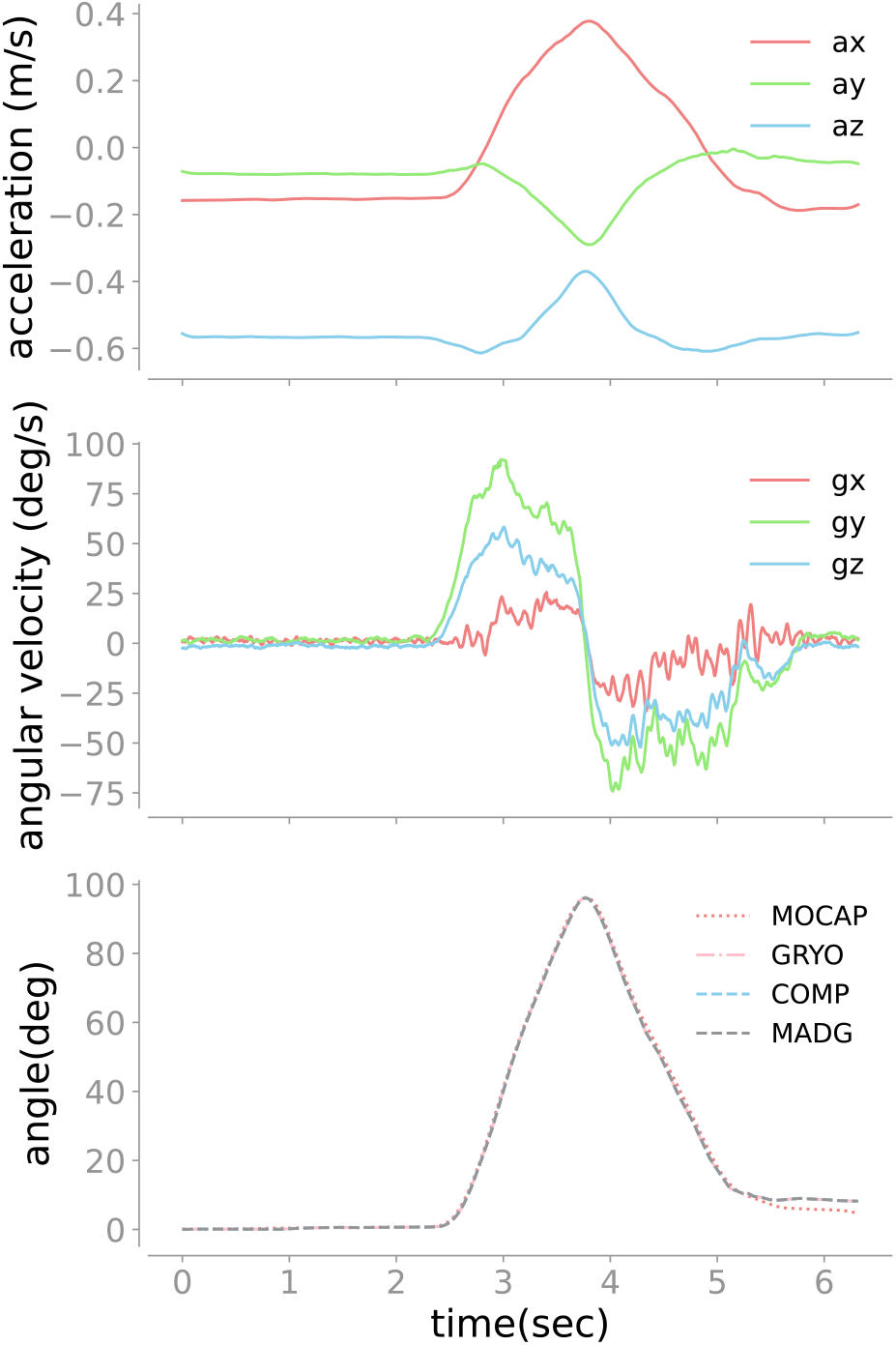
Raw data from the Romiumeter, The subplot on the top shows the acceleration along all 3 axes where acceleration along the x-axis is represented in light coral, along the y-axis is represented in light green, and along the z-axis is represented in sky blue. Similarly, the second subplot shows the angular velocity around all 3 axes with the same colour codes as the above plot for all axes. The subplot at the bottom shows the sample output of the angle reconstruction from the Romiumeter and OMCS. The reconstructed angles from the OMCS system are represented in light coral, while the reconstructed angle from the Romiumeter using the gyroscope integration, the Complementary filter, and the Madgwick filter are represented in pink, sky blue, and grey, respectively.

### 3.1 Angle reconstruction and ROM errors

Tables 1 and 2 present the reconstruction errors for the shoulder and neck movements from all three reconstruction methods – gyroscope integration, complementary filter, and the Madgwick filter. For both the shoulder and neck movements, the average reconstruction error (across all subjects and trials) was between 2.0 *−* 5.0^*°*^ across all three reconstruction methods. Interestingly, no significant differences were observed in the errors between the algorithms for both neck and shoulder movements (*p >* 0.9 for all movements). In fact, the average errors for neck and shoulder movements, as estimated by the gyroscope and complementary filter, were identical. This can be attributed to the near-zero optimal gain of the complementary filter, which means that the filter estimates most relied on the gyroscope integration.

**TABLE 1.**
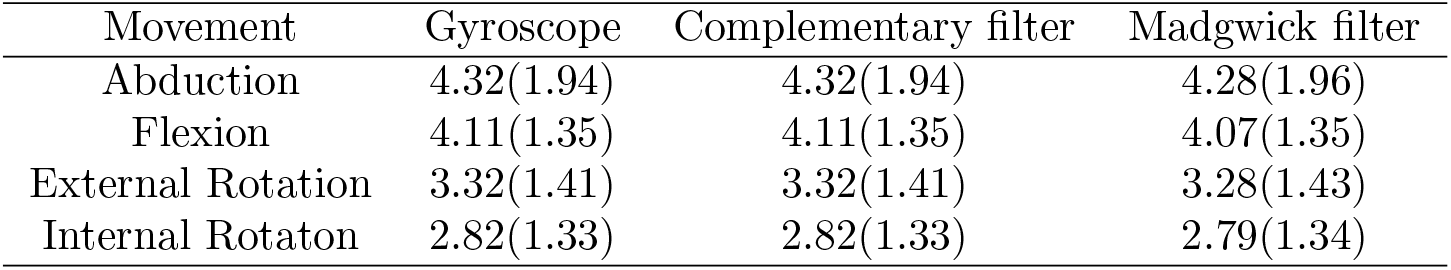
Maximum absolute error (degrees) of shoulder angles measured using Romiumeter and Motion capture system.

**TABLE 2.** Maximum absolute error (degrees) of neck angles measured using Romiumeter and Motion capture system.

| Movement | Gyroscope | Complementary filter | Madgwick filter |
| --- | --- | --- | --- |
| Flexion | 3.15(1.29) | 3.15(1.29) | 3.13(1.31) |
| Extension | 2.95(1.53) | 2.95(1.53) | 2.94(1.54) |
| Right Lateral Flexion | 3.10(2.66) | 3.10(2.66) | 3.08(2.66) |
| Left Lateral Flexion | 2.89(1.79) | 2.89(1.79) | 2.87(1.80) |
| Right Rotation | 4.68(2.60) | 4.68(2.60) | 4.63(2.64) |
| Left Rotation | 3.42(1.96) | 3.42(1.96) | 3.35(1.97) |

Given that no significant differences were observed between the algorithms, further analysis was conducted using the estimates from the gyroscope integration approach due to its simplicity. The average ROM errors for shoulder movements varied from 1.01^*°*^ (for internal rotation) to 2.40^*°*^ (for flexion/extension). Similarly, the average ROM errors for neck movements varied from 1.27^*°*^ (for left lateral flexion) to 2.19^*°*^ (for right rotation), as depicted in (Fig. 5).

**FGURE 5.**
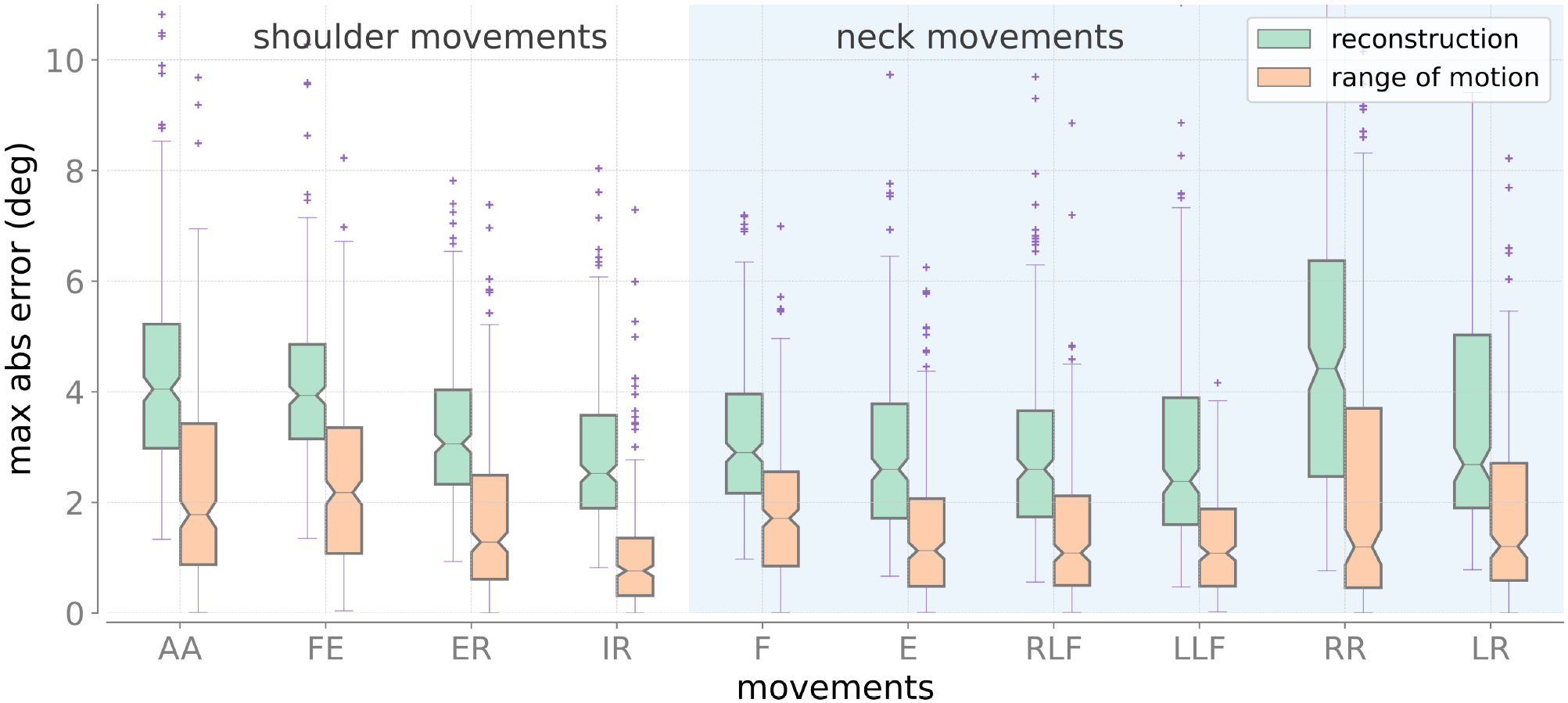
Reconstruction and range of motion errors estimated using gyroscope for shoulder and neck movements. AA: Abduction/Adduction, FE: Flexion/Extension, ER: External Rotation, IR: Internal Rotation, F: Flexion, E: Extension, RLF: Right Lateral Flexion, LLF: Left Lateral Flexion, RR: Right Rotation, LR: Left Rotation.

### 3.2 Intra-rater and inter-rater reliability

Tables 3 and 4 represent the intra-rater and inter-rater measurements of shoulder and neck ROM computed using gyroscope integration. For both shoulder and neck movements, the Romiumeter exhibited good reliability, ranging from 0.75 to 0.87. However, the inter-rater reliability of shoulder flexion (0.69) and right/left lateral neck flexion movements (0.72, 0.71) only showed moderate reliability.

**TABLE 3.** Intra-rater and Inter-rater reliability (ICC[95% CI]) of shoulder joint angle measurements using Romiumeter.

| Movement | Intra-rater | Inter-rater |
| --- | --- | --- |
| Abduction | 0.79 [0.59, 0.90] | 0.79 [0.58, 0.90] |
| Flexion | 0.76 [0.51, 0.88] | 0.69 [0.40, 0.85] |
| External Rotation | 0.87 [0.75, 0.94] | 0.75 [0.49, 0.88] |
| Internal Rotation | 0.76 [0.51, 0.88] | 0.77 [0.55, 0.89] |

**TABLE 4.**
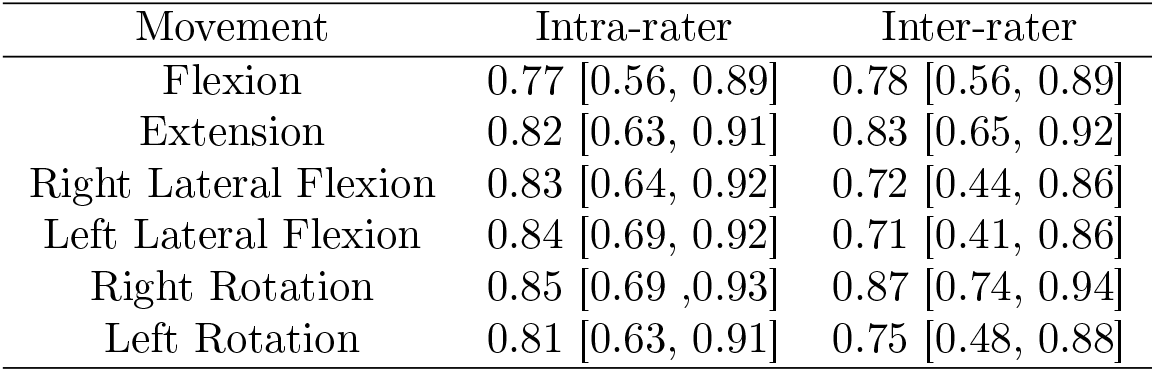
Intra-rater and Inter-rater reliability (ICC[95% CI]) of neck joint angle measurements using Romiumeter.

### 3.3 Sources of Reconstruction errors

Two additional experiments were carried out to understand the potential sources of the error in the angular reconstruction and the ROM estimates.

- **Effect of inherent gyroscope noise:** Fig. 6a represents the plot of the maximum absolute reconstruction error (in degrees) of the Romiumeter at rest for 10 sec computed using the gyroscope integration. The mean reconstruction error due to the inherent noise present in the gyroscope reached around 0.09*°* by 10s.
- **Effect of OMCS noise:** Fig. 6c shows the boxplot of the maximum absolute reconstruction (in green) and ROM (in orange) errors of the Romiumeter with respect to the encoder and OMCS, respectively. When compared with the high-resolution encoder, the mean reconstruction error from the gyroscope was less than 1.4∘. Whereas, when compared with the OMCS, the mean reconstruction errors were much higher.

**FGURE 6.**
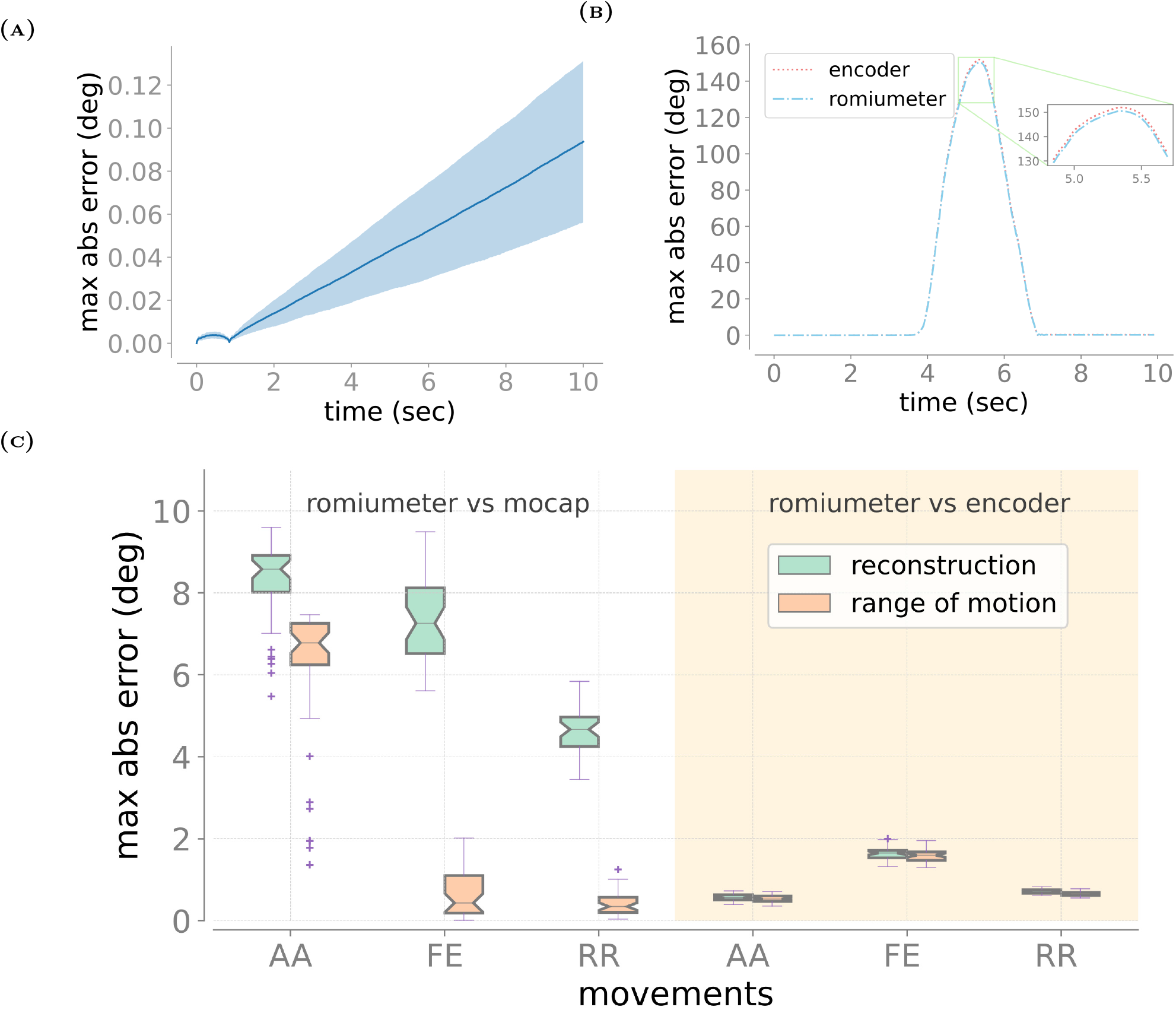
A) Reconstruction errors are estimated using the gyroscope when the sensor rests for 10 seconds. The blue line in the middle represents the mean of the data at each time point, and the shaded region extends up to one time of the standard deviation of the data at each time point. (B) Sample output of the reconstruction angle from the Romiumeter and angles measured by an encoder. (C) Reconstruction and range of motion errors were estimated using Romiumeter against OMCS and encoder in the additional experiment. AA: Abduction/Adduction, FE: Flexion/Extension, ER: External Rotation, IR: Internal Rotation, F: Flexion, E: Extension, RLF: Right Lateral Flexion, LLF: Left Lateral Flexion, RR: Right Rotation, LR: Left Rotation.

These experiments indicate that most reconstruction and ROM estimation errors might have been due to the noise in OMCS ground truth data.

## 4 Discussion

The present study showcased the validation of the Romiumeter against an OMCS system for measuring the range of motion in the shoulder and neck joints of asymptomatic individuals. The study compared various algorithms for reconstructing the angular kinematics from the accelerometer and gyroscope data. Additionally, it confirmed the reliability of measurements taken with the Romiumeter in asymptomatic individuals.

### 4.1 Validity of the Romiumeter

Compared with the OMCS, the Romiumeter demonstrated good accuracy, with a maximum absolute error of less than 5*°*. The majority of the studies found in the literature regarding ROM assessment of shoulder or neck movements using IMU have either used the root mean square error (RMSE), mean bias error (MBE), or mean absolute error (MAE) as a measure for validation [37–47]. We also computed RMSE, MBE, and MAE to compare our results with those of these studies. The average RMSE, MBE, and MAE for both shoulder and neck movements were less than 1.94*°*, 0.90*°*, and 1.42*°*, respectively. The MSE, MBE, and MAE values were lower in the current study [37–47]. Furthermore, most of these studies employed more than one IMU to measure ROM, while the current study employed a single IMU. This approach simplifies the assessment process for clinicians and requires patients to don/doff fewer devices. While Chan et al. and El-Gohary et al. assessed the ROM of the neck and shoulder, respectively, using a single IMU, the MBE was considerably higher in these studies (MBE > 5*°*) [41, 44]. Moreover, the Chan et al. study for measuring shoulder ROM required accurate alignment of the IMU’s axis of rotation with the anatomical axes of the humerus, making this method sensitive to a clinician’s expertise [41].

The gyroscope integration approach for ROM assessment was performed comparably with the Madgwick and the complementary filters for the current assessment protocol. The gyroscope integration approach was preferred due to its simplicity. The primary reason for the similar performances of the three methods could be the short-duration movements employed in the assessment protocol.

### 4.2 Reliability of Romiumeter

Intra-rater and inter-rater reliability were found to be good for all movements of shoulder and neck joints (ICC > 0.75), except that the inter-rater reliability (ICC: 0.69 - 0.72) of shoulder flexion, neck right lateral flexion, and left lateral flexion was moderate. The reason for these slight differences is unclear. These results are consistent with those of studies published by Anoro-Hervera et al. on intra-rater and inter-rater reliability of cervical active ROM in young asymptomatic adults (inter-reliability > 0.75 and intra-reliability > 0.90) [48] and Kaszyński et al. on the reliability of shoulder ROM using IMU sensors (inter-reliability: 0.88 - 0.97 and intra-reliability: 0.7 - 0.84) [45]. One important factor that a clinician needs to be aware of is the position of the joint before the movement because this can influence the reliability of the ROM assessment. If a participant’s starting joint position differs from their positions in previous sessions, it may lead to either overestimation or underestimation of ROM.

### 4.3 Angle Reconstruction and the Sources of Errors

Three algorithms were tested in the current study to reconstruct angular kinematics – gyroscope integration, the Complementary filter, and the Madgwick filter. To the best of our knowledge, this is the first study validating the shoulder/neck ROM assessment using IMU for different algorithms. No significant differences were found between these algorithms in measuring ROM, probably due to the relatively short duration of the movement trials employed in the current study. The simplicity of the gyroscope integration algorithm was the reason it was chosen for measuring ROM in the current study.

The maximum absolute error due to gyroscope integration drift when there is no movement is less than 0.2*°* for a 10-second-long integration duration (Figure 7(A)). When there is movement, the error is less than 2*°* (Figure 7(C)) compared with that of a high-resolution optical encoder. The errors are much larger when the same data was compared against the OMCS data (Figure 7(C)). The small reconstruction error when compared with that of the high-resolution encoder is similar to that of the study by Martyna et al. [43], who compared IMU-based angle reconstruction using a highly precise lightweight industrial robot (KUKA KR3 R540 AGILUS). They observed a RMSE of 0.15*°*, which was similar to the RMSE of 0.61*°* observed in the current study. Overall, these results indicate poor offset correction and errors in the OMCS ground truth data as the major sources of error in the current study.

**FGURE 7.**
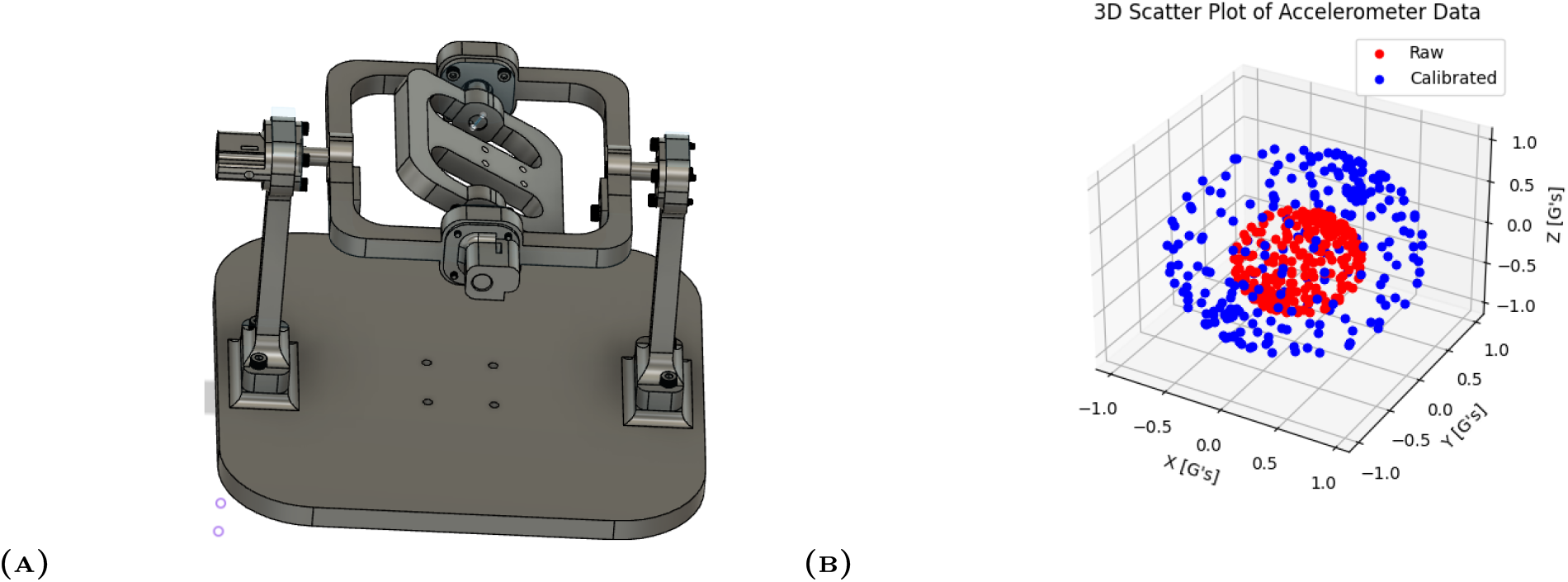
A) 3D model of the gimbal set-up used to calibrate the accelerometer. B) 3D scatter plot of raw accelerometer data and calibrated data. The blue dot represents the calibrated data and the red dot represents the raw data from the accelerometer

### 4.4 Limitations

The OMCS relied on only three markers to define the rigid body. Using additional markers could have reduced noise in the ground truth, potentially minimizing errors between the Romiumeter and the OMCS. Additionally, increasing the distance between markers might have improved the Romiumeter’s accuracy against the OMCS. The experimental group consists solely of asymptomatic individuals with no history of shoulder and neck pain in the past 3 months. However, it is important to note that this may not directly apply to symptomatic individuals. Future research should be aimed at exploring the test-retest reliability specifically in symptomatic individuals.

## 5. Conclusion

The paper introduces the Romiumeter, a single IMU-based device designed to measure the ROM of neck and shoulder movements. The Romiumeter demonstrated excellent accuracy in measuring ROM for both neck and shoulder movements. No significant differences were observed between the algorithms used for angle reconstruction, and gyroscope integration was chosen for further analysis due to its simplicity. Additionally, the Romiumeter exhibited moderate to good reliability in measuring the ROM for shoulder and neck movements. Future work must be aimed at evaluating the reliability of the Romiumeter in the patient population and exploring its usefulness for other joints as well.

## A. Appendix

### A.1 Details of the communication protocol

Each data packet consists of the following segments in the described order:

- **Start Bytes**: Two start bytes each valued 0xFF.
- **Size Byte**: A single byte indicates the payload size; this allows variable payload size for future expansion of the Romiumeter to include additional sensors.
- **IMU Data**: Six-byte long IMU data consisting of three accelerometer components followed by the three gyroscope components.
- **Sync Byte**: A single byte indicating the state of one of the Romiumeter’s digital pins to synchronize it to external triggers. The least significant bit is set to zero or one depending on the state of the digital pin.
- **Checksum**: A checksum byte which is equal to the sum of all the bytes from the start bytes to the sync byte is used to ensure packet integrity.

### A.2 Static calibration of the Romiumeter accelerometer

An accelerometer can be statically calibrated using gravity by varying the orientation of the accelerometer without imposing linear accelerations on the sensor. In this case, the magnitude of the acceleration measured by the accelerometer will be constant, equal to earth’s acceleration due to gravity. The purpose of the static calibration is to correct the non-orthogonality of the three sensing axes and to determine their corresponding scaling factors. The “raw” acceleration **a**_*raw*_ ∈ ℝ^3^ values recorded by the accelerometer and its relationship to the “true” acceleration **a**_*true*_ ∈ ℝ^3^ can be represented through the following model,

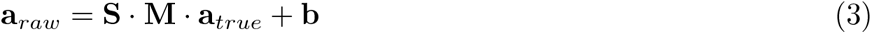

where, **S** ∈ ℝ^3*×*3^ is the sensitivity matrix, **M** ∈ ℝ^3*×*3^ is the matrix that accounts for the non-orthogonality of the three measurement directions of the accelerometer, and **b** ∈ ℝ^3^ is the bias vector. This calibration procedure will provide an estimate of the matrices **S** and **M**, along with bias vector **b**, which can be used to estimate the true value of the acceleration by inverting Eq. 3.

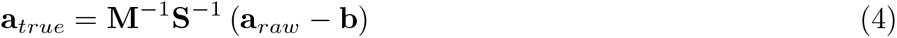

Renaudin et al. [49] presented an algorithm to calibrate a magnetometer based on the constraint that the norm of the Earth’s magnetic field is constant at a given geographical location. The same algorithm can be used to calibrate the accelerometer undergoing rotation without any linear acceleration. We designed a gimbal setup (Fig. 7a) to place the accelerometer in different orientations with respect to the earth’s gravitational field. We recorded data from the accelerometer by randomly placing it in arbitrary orientations. The raw accelerometer data were fit to an ellipsoid and the adaptive least squares estimator resulted in the required parameters for the calibration. Fig. 7b shows the 3D plot of the calibrated and uncalibrated data of the accelerometer.

### A.3 Details on gain optimization for Complementary Filter and Madgwick Filter

Gains for the Complementary filter and Madgwick filter were optimized using an approach similar to K-fold validation. For each movement, data was divided into 5 folds, and the average of the maximum absolute errors was computed for 4 folds and compared with the average of the maximum absolute error computed for the remaining one fold. This procedure was repeated till each fold is used as test data. The optimized gain was chosen based on which one resulted in less reconstruction error.

### A.4 Gyroscope Integration

#### Algorithm 1

Using the Gyroscope

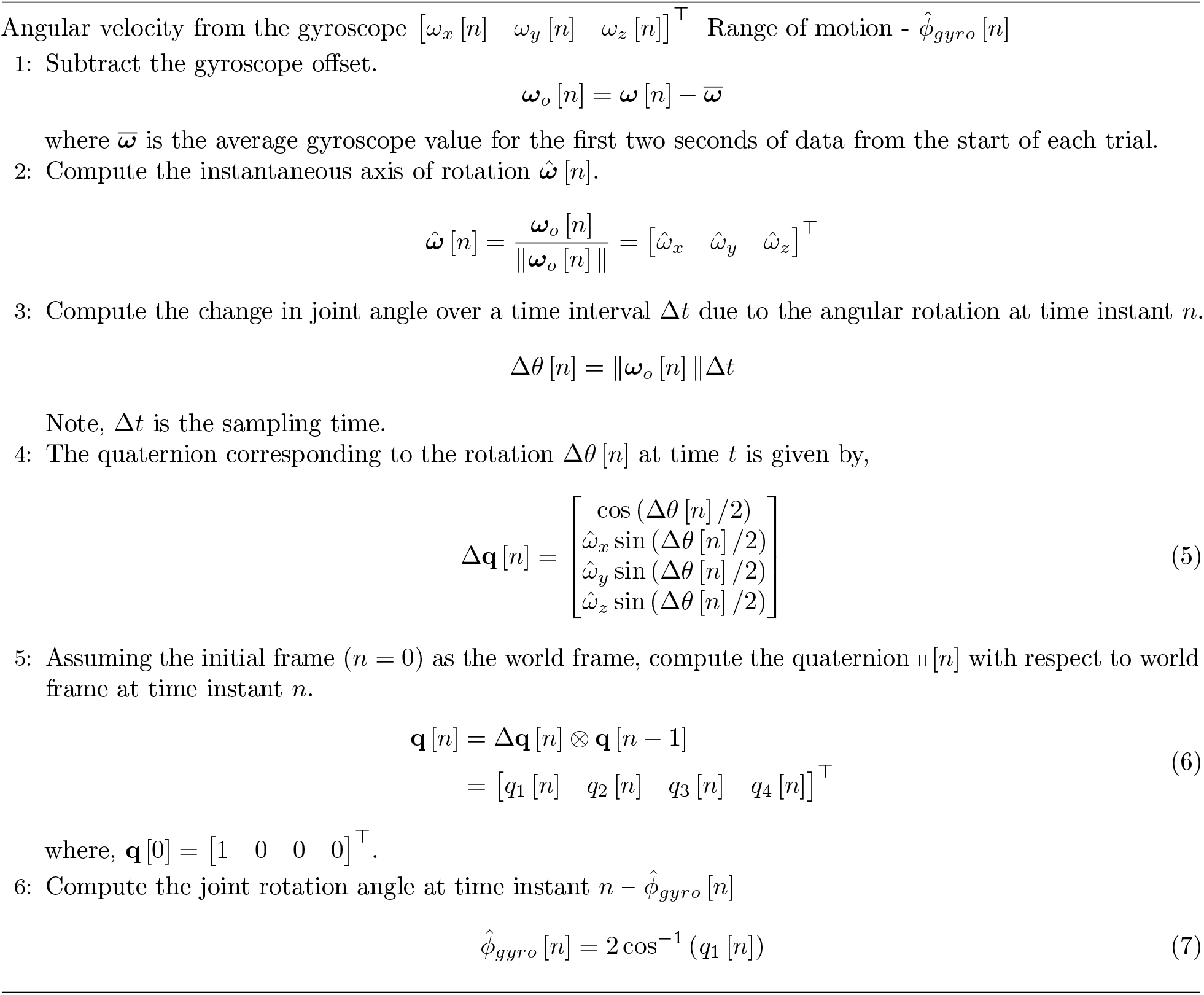

### A.5 Complementary Filter and Madgwick Filter

#### Algorithm 2

Complementary Filter

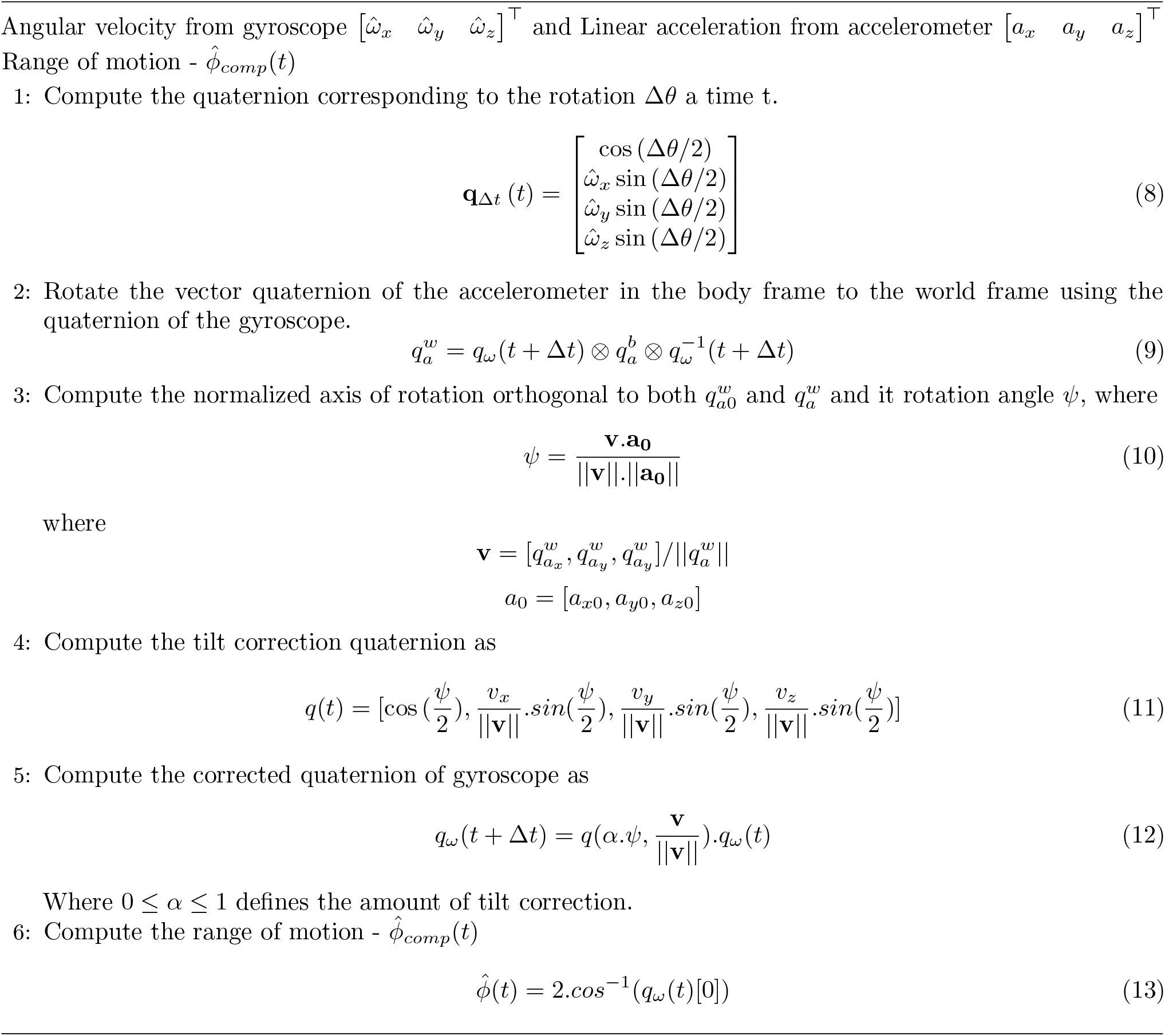

#### Algorithm 3

Madgwick Filter

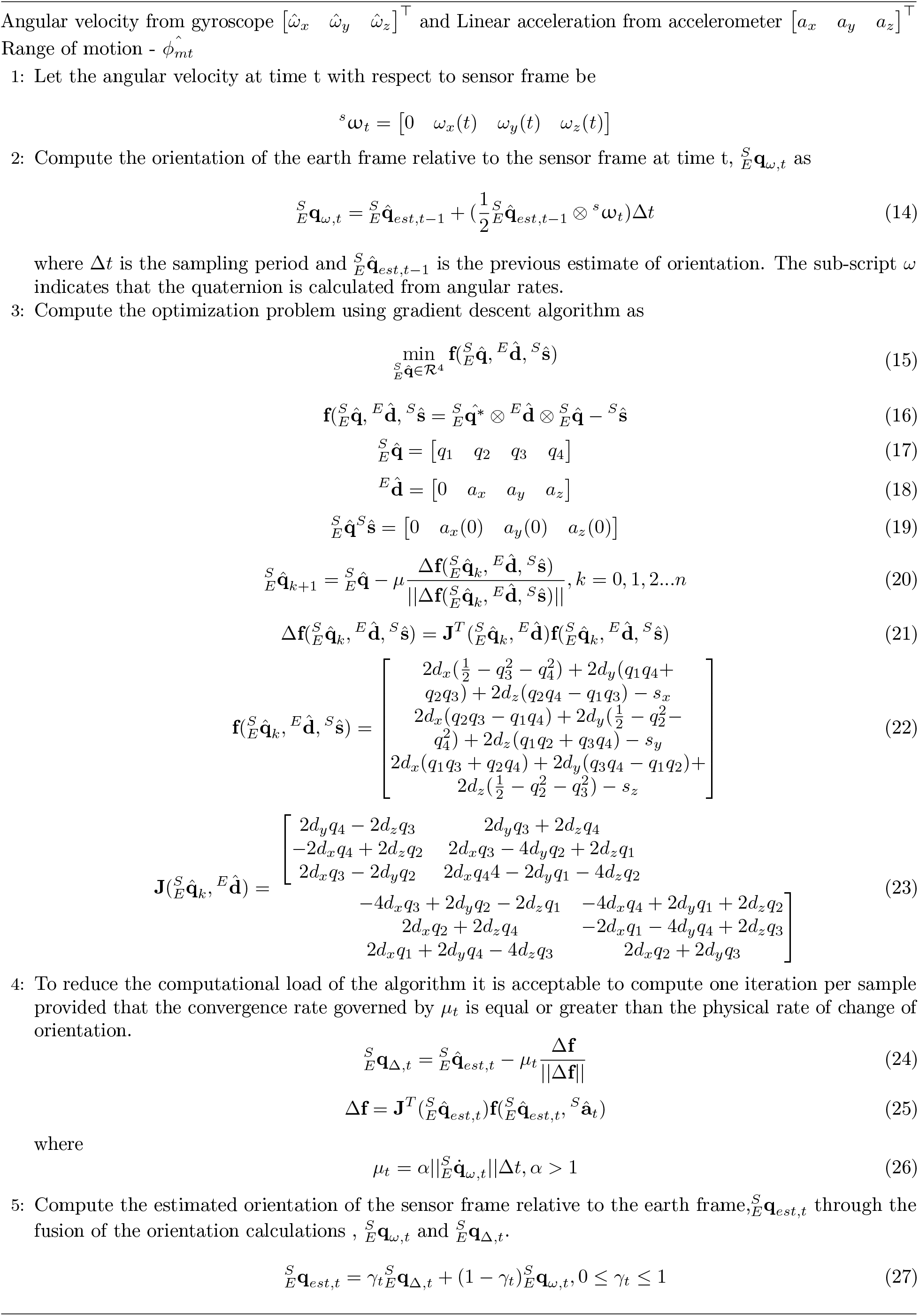

